# A Systematic Nomenclature for the *Drosophila* Ventral Nervous System

**DOI:** 10.1101/122952

**Authors:** Robert Court, Shigehiro Namiki, J Douglas Armstrong, Jana Börner, Gwyneth Card, Marta Costa, Michael Dickinson, Carsten Duch, Wyatt Korff, Richard Mann, David Merritt, Rod K Murphey, Andrew Seeds, Troy Shirangi, Julie H. Simpson, James W Truman, John Tuthill, Darren Williams, David Shepherd

## Abstract

The fruit fly, *Drosophila melanogaster*, is an established and powerful model system for neuroscience research with wide relevance in biology and medicine. Until recently, research on the *Drosophila* brain was hindered by the lack of a complete and uniform nomenclature. Recognising this problem, the Insect Brain Name Working Group produced an authoritative hierarchical nomenclature system for the adult insect brain, using *Drosophila melanogaster* as the reference framework, with other taxa considered to ensure greater consistency and expandability (Ito et al., 2014). Here, we extend this nomenclature system to the sub-gnathal regions of the adult *Drosophila* nervous system, thus providing a systematic anatomical description of the ventral nervous system (VNS). This portion of the nervous system includes the thoracic and abdominal neuromeres that were not included in the original work and contains the motor circuits that play essential roles in most fly behaviours.

## Background

Insects, and *Drosophila melanogaster* in particular, have made significant contributions to neuroscience research (Bellen et al., 2010). The powerful genetic tools and high-resolution neuroanatomy available in flies (Jenett et al., 2012; Scheffer and Meinertzhagen, 2019) and the large number of research groups working in this model, suggest that the rate of discovery will continue to be high. For the ventral nervous system, anatomical, physiological and molecular data are already available through the work of different laboratories. Integrating this information onto a common analytical framework is essential to provide an effective community-wide resource for future studies. Computational analysis combined with digital microscopy now make it possible to consolidate anatomical data from multiple techniques and transform how we analyse neural architecture and circuits (Jenett et al., 2006; Dance, 2015; Boettiger et al., 2016). The use of labelled 2D images to identify and define anatomical structures is no longer sufficient, as it is now possible to use multilayer microscopy with computational reconstruction to precisely define and allocate boundaries and structures in 3D. In addition, systems by which anatomical data gathered by different researchers may be registered on a common anatomical framework to provide a coarse understanding of connectivity are now available (Ostrovsky et al., 2013). Such methods rely on a systematic and consistent nomenclature that defines precisely the anatomical structures and their boundaries. Once this is complete it forms a coherent framework upon which information may be efficiently added, corrected, and extracted allowing new research findings to be added to the growing knowledgebase. Recognising this problem, a consortium of neurobiologists studying arthropod brains (the insect brain name working group (IBNWG), was established and produced a comprehensive hierarchical nomenclature system for the insect brain, using *Drosophila melanogaster* as the reference framework (Ito et al., 2014). This previous effort focused specifically on the brain and the gnathal regions that account for approximately 50% of the complete adult central nervous system. Here, we extend this nomenclature system to the sub-gnathal regions of the adult *Drosophila* nervous system; the region called the ventral nervous system (VNS). The VNS is the locus for the reception and integration of sensory information and also generates the locomotor actions that underlie most fly behaviors such as walking (Bidaye et al., 2014; Gowda et al., 2018; Mamiya et al., 2018; Mendes et al., 2013; Mendes et al., 2014; Tuthill and Wilson, 2016; Wosnitza et al., 2013), grooming (Seeds et al., 2014), jumping (Card and Dickinson, 2008), flying (Dickinson and Muijres, 2016), courtship (Clyne and Miesenböck, 2008; Shirangi et al., 2016) and copulation (Crickmore and Vosshall, 2013; Pavlou et al., 2016). Our work builds on previous anatomical descriptions of the ventral nervous system (Power, 1948; Merritt and Murphey, 1992; Boerner and Duch, 2010) and incorporates data from subsequent literature and findings made possible by new technologies and aims to resolve ambiguous terms. We propose a comprehensive and consistent nomenclature that will serve as a foundation for future work.

## Methodology

### Organization of the Working Group

The initial phase of work followed a similar format to that adopted by the original Insect Brain Name Working Group (IBNWG) to create the nomenclature for the *Drosophila* brain (Costa et al., 2013; Ito et al., 2014). We gathered researchers with expertise in the anatomy, development, and physiology of the VNS, hereafter referred to as the Drosophila Anatomy of the Ventral nervous system Working Group (DAVWG) for a workshop at the Janelia Research Campus in October 2013. We discussed a document listing all of the named regions found in the published literature and from the existing *Drosophila* anatomy ontology (Costa et al., 2013), as well as representative anatomical images assembled by authors Court and Shepherd. After systematic review and debate, the participants compiled a working proposal for wider comment. Iterative revisions resulted in the current nomenclature described here.

### Establishing the anatomical framework

Establishment of a systematic nomenclature requires a clear morphological and spatial definition of all the structures to be named and a standard naming scheme. The neuropil regions of the VNS are typically regarded as being ‘unstructured’ or ‘tangled’, or having a fine, granular appearance in sections with different regions distinguished only by general spatial terms such as, ‘intermediate’ (Merritt and Murphey, 1992) or ‘dorsal’. Despite this, different volumes of VNS neuropil can be defined in relation to fixed landmarks such as the longitudinal tracts and commissures (Shepherd et al., 2016).

Developmental origin provides an alternative organizational principle for defining the substructure of the neuropil. Neurons arise from neuroblasts whose first division results in A and B daughter cells. These undergo self-renewing divisions to produce clonal populations referred to a hemilineages. The neurons from a hemilineage tend to share properties, such as neurotransmitter identity and projection pattern – and even function (Harris et al., 2015; Lacin et al., 2019; Shepherd et al., 2019). Recently, Shepherd et al. (2016) used the primary projections of neuronal hemilineages to provide an organizational principle for defining the substructure of the neuropil. Although these landmarks may not always correspond to the underlying functional organization, they provide a consistent means of structurally defining neuropil regions.

To provide an initial framework for establishing distinct boundaries within the VNS, we generated confocal datasets that reveal various salient features, including tracts and neuropil. The anti-neuroglian antibody (Iwai et al., 1997) (Figure 1A-C) was used to reveal the projections of clonally related neurons in neuroblast (NB) hemilineages (Shepherd et al., 2016). An anti-Drosophila N-cadherin antibody (DNx8) was used to visualize neuropils according to the density of an active-zone-specific protein (Figure 1D and E) (Shepherd et al. 2016), mirroring exactly the structures revealed by nc82 (bruchpilot) immunostaining (Wagh et al., 2006). This allowed us to distinguish between neuropils that are poor in synapses, such as regions occupied by axons, primary neurites, and glial processes and synapse-rich regions such as the primary sensory neuropils and the dorsal neuropils associated with the neck, wings and halteres (Figure 1D and E). An anti-alpha tubulin antibody (data not shown) was used to reveal fibrous structures such as longitudinal tracts and commissures (Boerner and Duch, 2010). The images obtained with these labeling methods are available on the Virtual Fly Brain (https://github.com/VirtualFlyBrain/DrosAdultVNSdomains/tree/master/Court2017/template). Since all of these antibodies are available at low cost through the Developmental Studies Hybridoma Bank created by the NICHD of the NIH and maintained at The University of Iowa, Department of Biology, Iowa City, IA 52242, they can be used by future researchers to counterstain their own samples, identify neuropil regions described in this nomenclature, and computationally register them to our standard reference brains.

**Figure 1.**
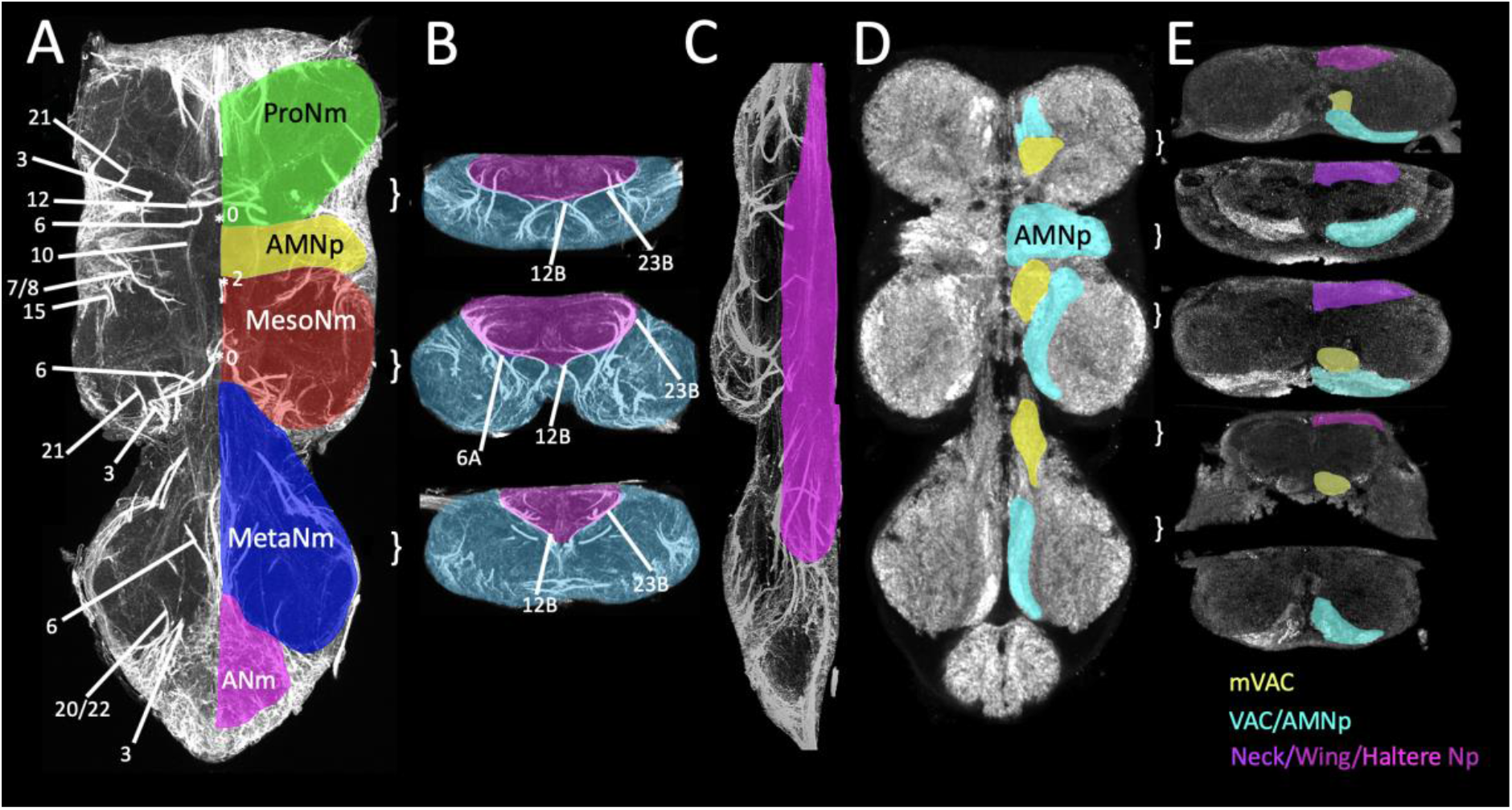
Selected sections through an adult VNS illustrating the tools used to define the major structures of the VNS. A-C) Neuroglian expression shown in A) horizontal, B) transverse and C) sagittal sections to reveal the tracts of the primary neurites of the postembryonic neuronal lineages. The pattern of labelled pathways is highly stereotyped, each pathway corresponds to the primary neurites of neurons derived from a single neuroblast. These tracts provide a robust basis for the identifying key structures of the VNS such as A) neuromere boundaries (ProNm, MesoNm, MetaNm and ANm). B and C) the ventral and anterior/posterior limits of the tectulum. D and E) General staining of the neuropil with N-cadherin, shown in D) horizontal and E) transverse sections, reveals the fine structure of the neuropil and regions of neuropil with high density staining. The regions of intense staining can be used to define and segment the VAC, mVAC, AMNp, neck neuropil, wing neuropil and haltere neuropil.

### The naming scheme

All of the anatomical data used in this manuscript can be found on the Virtual Flybrain website https://github.com/VirtualFlyBrain/DrosAdultVNSdomains. All of the text definitions of the structures considered in the nomenclature can be found on FlyBase: http://purl.obolibrary.org/obo/fbbt A key principle was to integrate existing terminology into the standard nomenclature we propose here. We made changes only to remove ambiguity. When multiple names for an anatomical entity were used in the literature, we gave preference to the name that was most commonly used based on citations. While we sought to preserve consistency with terms used for earlier developmental stages and in other insects, we avoided the implication of homology. Most of the naming scheme relies on morphological features rather than functional data, which we incorporate in the definitions when known. We also include a look-up table of synonyms, prior terms, and references.

### Abbreviations

We adopted a systematic approach when developing abbreviations for each named anatomical entity based on the following principles. (1) We adopted abbreviations that are unique across the whole CNS, avoiding abbreviations already in use for regions in the brain. (2) We created a system in which related entities would be easily recognizable. (3) We tried to be consistent with nomenclature established for the brain (Ito et al., 2014). The reasoning behind each abbreviation change was recorded and embedded in the definition. When referring to the neuromere and related structures, abbreviations were changed from a single letter or number to ‘Pro’, ‘Meso’ and ‘Meta’. This removed confusion with positional abbreviations such as posterior or medial. The use of the single letter ‘N’, which is used widely (neuromere, neuropil, nerve, neuron), was reserved for “nerve”; other larger gross anatomy structures differentiated with additional letters (e.g. ‘Nm’ for neuromere and ‘Np’ for neuropil). The letter ‘C’ was used to identify commissures. In cases where multiple abbreviations already exist in the literature for specific structures, the abbreviation that provided the clearest indication with least likelihood of confusion was selected and additional abbreviations were captured as synonyms.

### Axis orientation

The general axis of orientation for the VNS is straightforward. The neuroaxis and the body axis are the same, with the prothoracic neuromere being anterior most and the abdomen (abdominal ganglionic complex) being the most posterior. In the dorsal/ventral plane, the tectulum is dorsal and the leg nerves ventral. The dorsal/ventral axis is also sometimes referred to as superior/inferior but dorsal and ventral are the preferred terms. The designation of left and right is assigned as if the sample is viewed from above (dorsal). The orientation in all figures is indicated by arrows pointing towards the anterior (A), dorsal (D) and the right (R).

### Definition of the VNS

The VNS is the region of the central nervous system posterior to the brain. It is connected to the brain by descending and ascending neurons that pass through the neck connective. The *Drosophila* VNS is a single consolidated ganglion located in the ventral part of the thorax. This ganglion contains all of the thoracic and abdominal neuromeres (Figure 1) and was called the thoracicoabdominal ganglion by Power (1948). The VNS is also often referred to as the ventral nerve cord (VNC), and we keep this term as an acceptable synonym. We prefer the term VNS because it explicitly excludes the gnathal neuromeres (Niven et al., 2008), those associated with the proboscis, which have been included as the Suboesophageal Zone in the brain nomenclature.

### Identifying and defining the neuropil structures in the VNS

Many insects have a ladder-like ventral nervous system composed of physically separated segmental neuromeres connected by longitudinal tracts, but in *Drosophila*, the thoracic and abdominal neuromeres are fused into a single complex (Niven et al., 2008). At the gross anatomical level, the segmental organization of the VNS can be resolved from external morphology. The thoracic neuromeres constitute the bulk of the VNS and are recognisable as three paired enlargements at the anterior of the VNS, corresponding to the prothoracic, mesothoracic and metathoracic neuromeres (ProNm, MesoNm and MetaNm, Figure 1A). At the posterior end is a small, dorsally located mass, the abdominal neuromeres, that is a fusion of all the abdominal neuromeres (ANm, Figure 1A). Despite the evident external segmental organization of the VNS the fusion of multiple neuromeres means that identifying precise neuropil boundaries can be problematic. One of our aims was to define different regions of neuropil and provide landmarks to facilitate consistent identifications and nomenclature for future studies. Although the VNS does not have the clearly defined compartmental structure found in the *Drosophila* brain it does have a clear architecture of tracts, commissures and primary axon bundles that provide the basis for defining different regions of neuropil. Cell body positions are not a reliable indicator of the segmental organization of the VNS. There are many examples of cell bodies being passively displaced during neuropil expansion at metamorphosis resulting in somata being drawn across the midline or pulled into adjacent neuromeres (Shepherd et al., 2019).

## Major neuropil features of the VNS

### Neuromere boundaries

Although the VNS is a fusion of thoracic and abdominal neuromeres, it is possible to define the boundaries of the major neuromeres using the scaffold of axon fibres revealed by neuroglian expression. The neuroglian positive axonal bundles are formed from tightly fasciculated primary axons from individual neuronal lineages. Since each neuromere is founded by a specific set of NBs, the neuroglian bundles create a neuromere specific set of anatomical markers that provide a robust framework to define each neuromere (Figure 1A-C) (Shepherd et al, 2016).

**The Prothoracic Neuromere** (ProNm; also called T1) is the anterior-most VNS neuromere and its anterior boundary defines the anterior extent of the VNS. Although the posterior boundary is less obvious, it can be defined by the extent of the primary neurite projections of the central neurons produced by the prothoracic NBs (0, 3, 6, 11, 19 and 21). These neuroglian positive tracts project anteriorly into the neuromere (Figure 1A) and their entry points represent the posterior limit of the prothoracic neuropil.

**The Mesothoracic Neuromere** (MesoNm; also called T2) is delimited anteriorly by the entry points of the neuroglian-positive tracts from the anterior mesothoracic lineages 2, 7, 8, 10, 15 and 16 (Figure 1A). The posterior margin is defined by the entry points of the neuroglian-positive tracts from the posterior mesothoracic lineages 0, 3, 6, 11, 19, and 21, all of which project anteriorly into the neuromere (Figure 1A).

**The Metathoracic Neuromere** (MetaNm; also called T3) is defined anteriorly by the tracts of theposterior mesothoracic lineages 0, 3, 6, 11, 19, and 21 (see above and Figure 1A). Since morphogenetic movements draw the metathoracic cell bodies posteriorly, these lineages have extended primary neurites that do not clearly define the anterior border. The posterior margin of the metathoracic neuromere is best defined by the entry points of the primary projections from the posterior lineages 0, 3, 6, 19, 20/22 and 21, only some of which are visible in the plane shown in Figure 1A.

### Other Neuropils

The fusion of the thoracic neuromeres created two additional major neuropil regions, the accessory mesothoracic neuropil (AMNp) and the tectulum, that do not conform to the evolutionarily ancestral segmental origins.

**The accessory mesothoracic neuropil (AMNp)** is a subdivision of the mesothoracic neuromere located at the interface between the pro- and mesothoracic neuromeres. The AMNp is bounded anteriorly by the entry points of the posterior prothoracic lineages 0, 5, 6, 11, 19 and 23 and posteriorly by the entry points anterior mesothoracic lineages 1, 2, 7, 8, 9, 15, and 16 (Figure 1A) (Shepherd et al., 2016). The AMNp also contains a dense synaptic neuropil (as seen by brighter signal in NC82 or N-cadherin staining in Figure 1D and E) that is derived from sensory afferents from the wing and notum entering the VNS via the Anterior Dorsal Median Nerve (ADMN) (Power, 1948). This dense synaptic neuropil corresponds to a structure called the ovoid by Merritt and Murphey (1992). The AMNp was originally called the Accessory Mesothoracic Neuromere by Merritt and Murphey (1992) but we substitute *neuropil* to indicate that it is a region made up of incoming sensory afferents rather than intrinsic neurons of a common developmental origin.

**The tectulum** is specialized region of the dorsal VNS neuropil. Whilst the neuromeres divide the VNS along the anterior-to-posterior axis, there is also specialization on the dorso-ventral axis. The tectulum (Tct) was described by Power (1948) as a discrete dorsal region of the VNS, overlying the mesothoracic neuromere like a saddle, and extending over the posterior prothoracic and anterior metathoracic neuromeres. As with the neuromeres, the neuroglian positive primary neurites provide boundaries that precisely circumscribe the tectulum to define its boundaries (Figure 1B and C) (Shepherd et al., 2016). On this basis the tectulum was defined as a region of neuropil dorsal to hemilineage tracts 12B, 6A, and 23B in all three thoracic neuromeres that extends posteriorly from the cervical connectives in the prothorax through the mesothorax with its posterior most point defined by the entry point of lineage 3 in the metathorax (Figure 1B and C).

Although we originally, defined the tectulum as a single neuropil without sub-divisions a detailed analysis of the innervation patterns of descending interneurons to the VNS has suggested that the tectulum can be stratified into three layers: a ventral layer (the lower tectulum), an intermediate layer (the tectulum) and a dorsal layer (the upper tectulum) (Figures 1E, 4 and 5; Namiki et al., 2018). Whilst the identification of these three layers is anatomically correct the nomenclature used by Namiki et al. (2018) is problematic and required a revision.

Namiki et al. (2018) define the **lower tectulum** as a region of the pro and mesothoracic neuromeres sandwiched between the prothoracic mVAC and the posterior margin of the mesothoracic neuromere at the ITD-CFF commissure. It is contained ventrally by the tract VTV and laterally by DLV tract (Figure 5). Whilst we accept the identification of this region of neuropil the problem with Namiki’s definition is that the defined region does not fit with the accepted definition of the tectulum because the newly described lower tectulum is ventral to hemilineage projections (12B, 6A, and 23B, Figure 1A) and not part of Power’s original definition of the tectulum. To resolve this inconsistency, we propose that this newly defined neuropil be renamed the **Central Association Center** and not lower tectulum. The name is selected to reflect the previous use of the term Association Center to describe different neuropils in the VNS (mVAC and VAC) (Tyrer and Gregory, 1982; Merritt and Murphey, 1992).

The middle stratum, called the tectulum by Namiki et al. (2018), does not match with the accepted definition of the tectulum. It does not include the dorsal neuropils, which they segmented out as the newly named upper tectulum (see below). Whilst Namiki et al.’s (2018) description of this middle stratum in the tectulum is based on solid evidence, to avoid the confusion created by using the same name (tectulum) to describe different neuropils, we concluded that the neuropil called the tectulum by Namiki et al. (2018) should be renamed as **Intermediate Tectulum**.

The third stratum of the tectulum, called the upper tectulum by Namiki et al., (2018), is the dorsal most part of the tectulum, dorsal to the tracts DMT, IDT and ITD-HC. There are no issue with this nomenclature and upper tectulum is the accepted name for this neuropil. The upper tectulum can be further segregated on the basis of the enriched N-Cadherin/bruchpilot expression (Figure 1D and E) into three neuromere specific neuropils. A prothoracic neuropil called neck neuropil (Neck, Figures 1D and E, 4 and 5), a mesothoracic neuropil called wing neuropil (Wing, Figures 1D and E, 4 and 5) and a metathoracic neuropil called the haltere neuropil (Haltere, Figures 1D and E, 4 and 5).

#### The Leg Neuropil

If the tectulum represents the dorsal third of the thoracic neuromeres, the ventral two-thirds can be considered leg neuropils. These regions receive sensory inputs from the legs and house the motor neurons that target leg muscles, as well as many other local interneurons and projecting neurites. Unlike the tectulum, the leg neuropils exhibit clear segmental boundaries and although each thoracic neuromere is slightly different they all conform to the same organizational principles. The leg neuropils are best described in transverse section: The leg neuropil can be partitioned into distinct regions along the dorsoventral axis. The ventral-most layer of leg neuropil, the Ventral Association Center (VAC) (Merritt and Murphey, 1992) is readily distinguishable as a unique region by the high expression levels of synaptic antigens bruchpilot and N-cadherin (VAC, Figures 1D and E and 5) which are specific to synaptic components (Wagh et al., 2006) and (Iwai et al., 1997) indicating high synaptic density in VAC. The VAC is innervated by sensory afferents from sensory neurons associated with tactile bristles on the leg which form a somatotopic projection (Murphey et al., 1989) within the VAC. Adjacent to the VAC of each leg neuropil is a paired globular structure, the medial Ventral Association Center (mVAC), a bilaterally symmetrical region that can be identified both by its fine textured appearance and the high expression levels of bruchpilot and N-cadherin (mVAC, Figures 1D and E) (Merritt and Murphey, 1992). In *Drosophila* the mVAC is innervated by a subset of Femoral Chordotonal Organ (FCO) sensory neurons which form a “club” shaped projection that terminates in the mVAC (Phillis et al., 1996). The Drosophila mVAC is almost certainly homologous to the mVAC described in locusts and other insects which also receive primary sensory afferents for leg chordotonal organs and is known as “auditory neuropil” (Römer et al., 1988; Oshinsky and Hoy, 2002).

The leg neuropil, between the VAC and the tectulum, is called “intermediate neuropil” (IntNp) because it occupies most of the central third of the dorsoventral area in transverse section (IntNp, Figures 4 and 5). The leg neuropil contains the dendritic branches of the leg motorneurons as well premotor interneurons (Shepherd et al., 2019) and sensory afferent terminals from leg campaniform sensilla, hair plates and the “hook” and “claw” projection types from the FCO (Mamiya et al., 2018). Like the tectulum, the leg neuropils exhibit clear functional segregation: Motor neurons are located dorsally and the sensory modalities are partitioned into layers, with proprioception in intermediate neuropil, and a somatotopic representation of tactile information in the ventral-most zone (Murphey et al., 1989; Tsubouchi et al., 2017).

#### Abdominal Neuromeres (ANm)

The remaining major neuropil of the VNS is the abdominal neuropil (ANp), which is a fusion of abdominal neuromeres A1 through A8. The lineage composition of the abdominal neuromeres is not known and it is not possible to define the neuropils of individual neuromeres but the anterior limit of the ANm can be defined by reference to the neuroglian bundles that delineate the posterior limit of the MetaNm, lineages 0, 3, 6, 20/22 and 23 (Figure 1A).

### Tracts and Commissures

Building on classic studies of orthopterous insect ganglia such as the grasshopper (Tyrer and Gregory, 1982), Merritt and Murphey, (1992) and Boerner and Duch (2010) were able to describe in detail, the stereotyped patterns of longitudinal tracts and commissures in the adult Drosophila VNS (Figures 2, 3 and 5). These studies, along with the work of Shepherd et al. (2016) using postembryonic neuron lineages, define the majority of adult VNS commissures and tracts. These provide landmarks for describing locations in the VNS and we provide a summary of the consensus nomenclature here.

**Figure 2.**
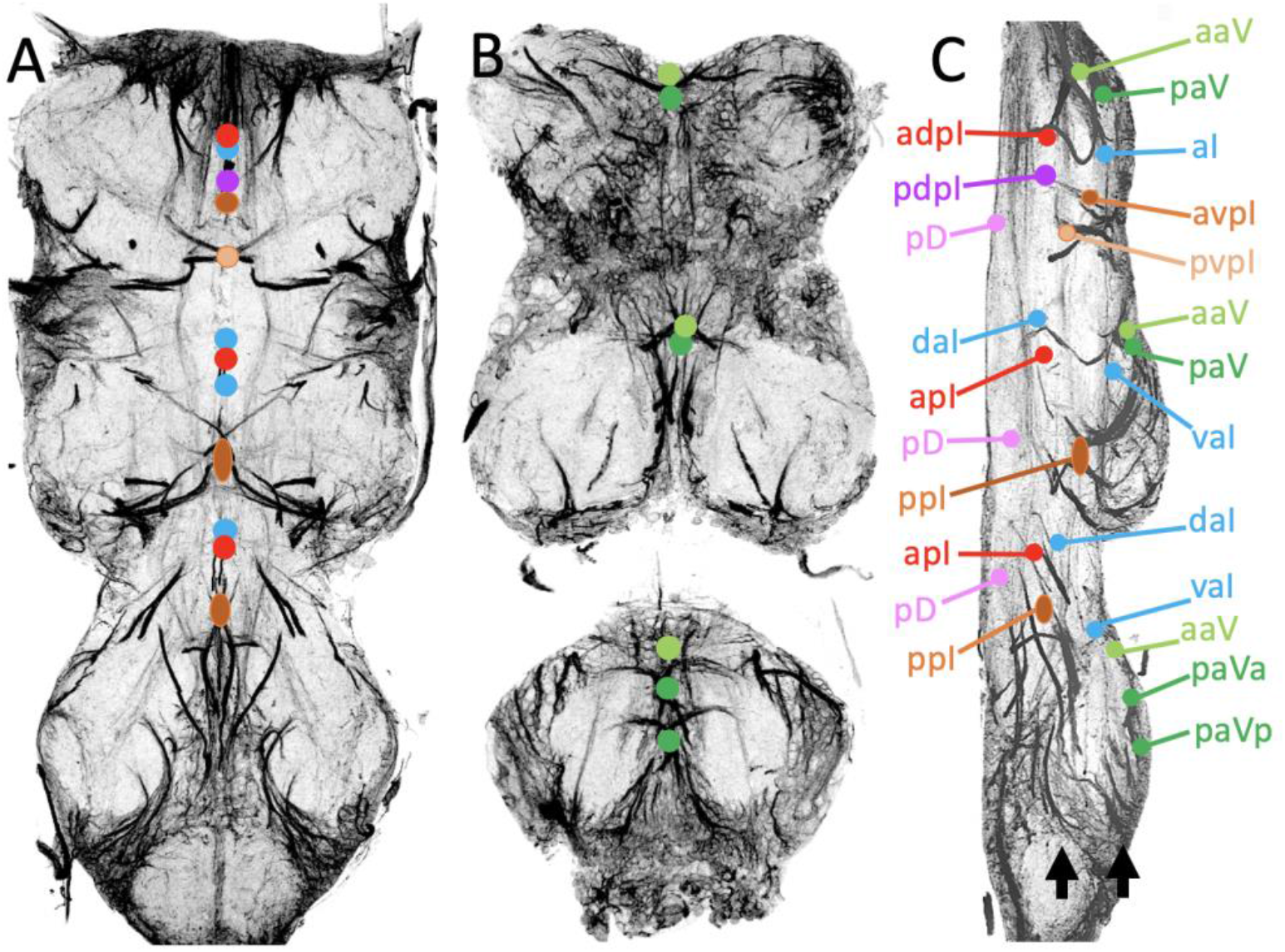
Identification of the major Commissural Pathways of the VNS. Neuroglian expression of the primary neurite projections of neuronal hemilineages can be used to identify the major commissural pathways in the VNS. A and B) Horizontal sections through the VNS with the major commissures indicated by colored dots. The plane of each section is shown by the black arrows in Panel C). C) Sagittal section of the VNS again showing the locations of the major commissural pathways.

**Figure 3.**
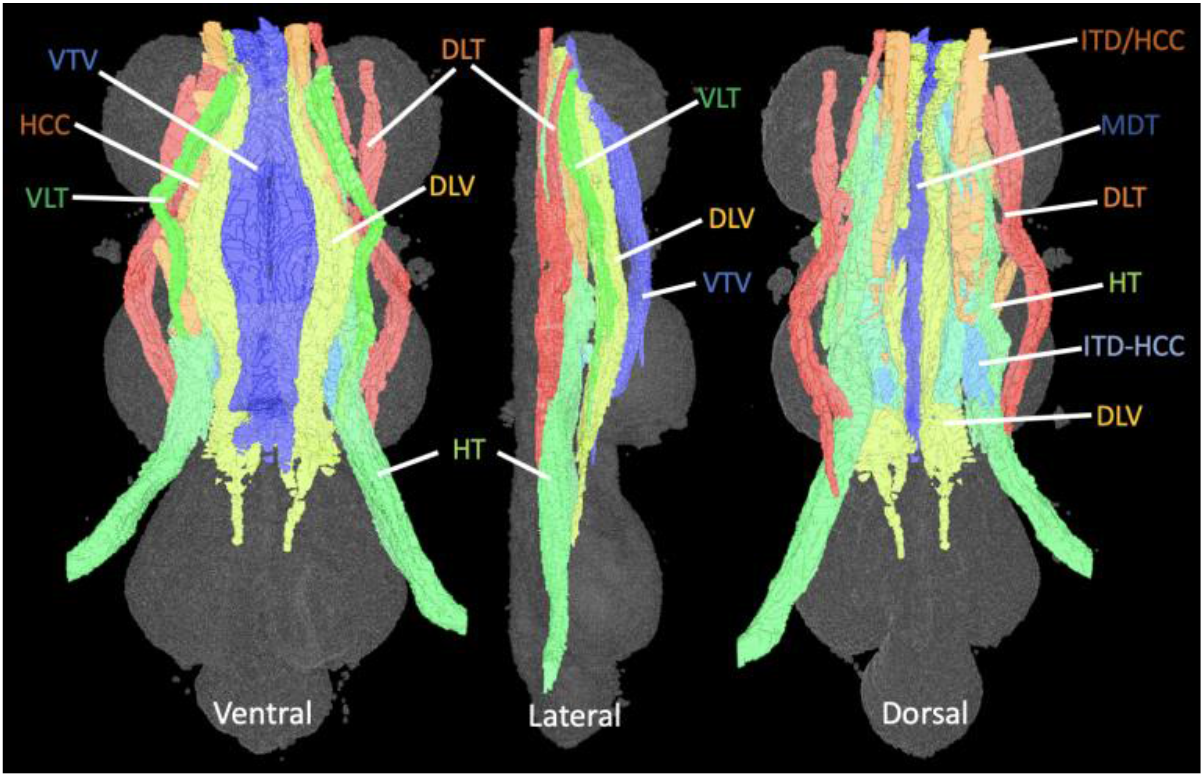
The Longitudinal Tracts. Surface rendering of the major tracts segmented from tubulin labelled VNS. The tracts are illustrated from ventral, lateral and dorsal perspectives (left to right).

### The Commissures

The larval VNS has five commissures per neuromere: the anterior (aD) and posterior (pD) dorsal commissures, the anterior (aI) and posterior (pI) intermediate commissures, and the anterior ventral (aV) commissure (Truman et al., 2004). Each postembryonic lineage that crosses the midline does so via a specific and invariant commissure (Truman et al., 2004). The five larval commissures segregate into additional pathways during metamorphosis, so the adult fly has many more commissures. Using lineage-based markers, Shepherd et al (2016) linked the larval commissures to their adult counterpart (Power, 1948; Merritt and Murphey, 1992). These lineage-based definitions underlie the proposed nomenclature.

#### Commissures derived from the larval aV commissure

The larval aV commissure is present in all three thoracic neuromeres. In the adult thoracic neuromeres it segregates into two distinct commissures which we called the **anterior anterior Ventral Commissure** (aaV) and the **posterior anterior Ventral Commissure** (paV) (Figure 2). The aaV is formed by the axons of hemilineage 1A and sits at the anterior of the neuromere at the ventral-most margins, outside the neuropil and cell cortex and anterior to the hemilineage 2A axons. The paV is formed by axons of hemilineages 13B and 14A and sits at the anterior of the ProNm but crosses the midline posterior to the hemilineage 2A primary neurites (Shepherd et al., 2016). In MetaNm the axons of paV are pulled apart to form two distinct commissures (paVa and paVp, Figure 2). Only two of these commissures were described by Power (1948): the MesoNm paV was called the Ventral Accessory Commissure of the Mesothoracic Neuromere by Power and the MetaNm aaV was called the Accessory Commissure of the Metathoracic Neuromere.

#### Commissures derived from the larval aI commissure

The larval anterior Intermediate Commissure (aI) is present in all three thoracic neuromeres. In the prothoracic neuromere it contains only 10B axons but in the meso and metathoracic neuromeres it contains the axons from hemilineages 10B and 18B. In the adult, the ProNm retains a single aI commissure; the **anterior Intermediate Commissure** (**aI**, Figure 2) but in the MesoNm and MetaNm aI segregates to form two commissures the **ventral anterior Intermediate Commissure (vaI)** formed by the 10B neurons (**vaI**, Figure 2) and the **dorsal anterior Intermediate Commissure** (**daI**) (**daI**, Figure 2) formed by the 18B neurons. In all 3 neuromeres these commissures are ventrally located at the anterior of the neuromere and posterior to the hemilineage 2A primary neurites (Shepherd et al., 2016).

#### Commissures derived from the larval pI commissure

The larval posterior Intermediate Commissure (pI) is formed by axons from hemilineages 5, 6B, 7, 8 and 12B. In adult MesoNm and MetaNm this commissure segregates into two commissures, one containing axons from hemilineages 7B and 8B and the other containing axons from 5, 6B and 12B. The anterior-most, containing the hemilineages 7B and 8B axons, creates the two most robust commissures of the adult VNS which Power (1948) called the **Commissure of the Mesothoracic Neuromere** (mesothorax) and the **Haltere Commissure** (metathorax). To harmonize the nomenclature, we renamed them both as the **anterior pI Commissure (apI)** (Figure 2). The commissure formed by the axons from hemilineage 5B, 6B and 12B was not previously identified and is called the **posterior pI Commissure (ppI)** (Figure 2).

In the prothoracic neuromere the organization is more complex as the larval pI segregates into 4 commissures. Hemilineage 7B and 8B axons separate to form two commissures and the hemilineage 5B axons separate from 6B and 12B (Figure 2). None of these commissures had been previously described. The commissure formed by hemilineage 7 is called the **anterior dorsal pI Commissure** (adpI), the commissure formed by hemilineage 8B is called the **posterior dorsal pi Commissure (pdpI),** the commissure formed by 5B is called the **anterior ventral pI Commissure (avpI)** and the commissure formed by 6B and 12B is called the **posterior ventral pI Commissure (pvpI).**

#### Commissures derived from the larval pD commissure

The larval posterior Dorsal commissure (pD) is the dorsal-most commissure and is found in all three thoracic neuromeres. It contains the axons from hemilineage 6A. In the adult the commissure is also the dorsal most and located in the upper tectulum. Power (1948) only described this commissure in the mesothorax and called it the Posterior Dorsal Mesothoracic Decussation. We called all three commissures the **Posterior Dorsal Commissures (pD).**

#### Commissures formed by descending neurons

In the mesothoracic neuromere there is a robust commissure formed by the inner tracts of the intermediate tract of the dorsal cervical fasciculus (ITD-CFF), as they cross the midline, anterior to the haltere commissure in the upper tectulum above the mesothoracic neuromere to terminate on the contralateral side. This was termed Commissure of the Fine Fibers of the Intermediate Tract of the Dorsal Cervical Fasciculus (CFF) by Power (1948.

#### Other Commissures

In addition to the commissures that can be related to the projections of specific hemilineages there are two commissures described by Power (1948) that cannot be related to specific hemilineages. **The Commissure of Prothoracic Neuromeres (CPN)** is transverse bundle of fibers that cross the midline in the ProNm. The fibers characteristically bow posteriorly and are dorsal to the dorsal lateral tracts of the ventral cervical fasciculus (DLV). The **Dorsal Accessory Commissure of the Mesothoracic Neuromeres (DAM)** Is a commissure in the dorsoposterior region of the MesoNm but ventral to the entry point of the dorsal metathoracic (haltere) nerves and ventroanterior to the **Mesothoracic pdpI Commissures.**

### Longitudinal Tracts

The commissural pathways described above contain neurites that cross the midline, carrying contralateral projections. In contrast, we use the term longitudinal tracts to describe the ordered and invariant array of eight bilaterally symmetrical axon bundles that run parallel to the anterior-posterior axis of the ventral nervous system (Figure 3). Unlike the commissures, the longitudinal tracts were fully described by Power (1948) and Merritt and Murphey (1992) with a largely consistent nomenclature which we summarize here.

#### Dorsal Lateral Tract (DLT)

As its name implies the DLT is located in dorsal lateral neuropil (Figure 3), it is formed by fibers from the lateral bundles in the cervical (neck) connective and projects posteriorly and superficially at the dorsal lateral edge of the neuropil to terminate in the metathoracic neuromere (Figure 3). The tract contains the axons of descending neurons that innervate neck, wing, haltere and leg neuropils (Namiki et al, 2018). Histologically DLT has coarser fibers than the other tracts derived from the ventral bundles of the cervical connective described below.

#### Intermediate Tract of Dorsal Cervical Fasciculus (ITD)

The ITD is a dorsal tract, derived from the dorsal fibers in the connective and sits just medial to the DLT (Figure 3). According to Power (1948), the ITD projects posteriorly and separates into three adjacent tracts. The medial most of these subdivisions, called ITD-CFF turns medially in the mesothoracic neuromere to cross its contralateral homolog to form the chiasma of fine fibers of the intermediate tracts of the dorsal cervical fasciculus (ITD-CFF). This longitudinal projection is what we now refer to as ITD. It contains the axons of many descending interneurons that terminate widely in neck, wing, haltere and leg neuropils (Namiki et al 2018). The other two subdivisions of ITD are now recognized as distinct tracts and called the Haltere Chiasma (ITD-HC) and the Haltere Tract (ITD-HT) respectively.

#### Intermediate Tract of Dorsal Cervical Fasciculus – Haltere Chiasma (ITD-HC)

ITD-HC is formed by the axons of the cHIN interneurons as they project anteriorly from the metathoracic neuromere. The axons originate from the interneurons produced by metathoracic hemilineage 8B, the primary projections of which also form the major component of the Haltere Commissure. The tract itself extends anteriorly from the HC just lateral to the ITD and medial to the HC (Figure 3 and 5). This tract was termed cHiN by Merritt and Murphey (1992).

#### Haltere Tract ITD-(HT)

The haltere tract is the most lateral component of Power’s (1948) ITD (Figure 4) and is composed of many large-diameter fibers that can be traced as a bundle into the cervical connective. The HT is formed by the sensory afferent axons (Ghysen, 1980; Strausfeld and Seyan, 1985) from the dorsal metathoracic nerve (Haltere Nerve) entering the metathoracic neuromere and extending anteriorly through the cervical connective (Power, 1948; Merritt and Murphey, 1992). The tract has small arborizations with some of the Fibers bending anterolaterally to become part of the haltere commissure (HC) in the metathoracic neuromere, while others turn ventrally and straggle into the dorsolateral part of the mesothoracic neuromere where they are quickly lost (Power, 1948).

**Figure 4.**
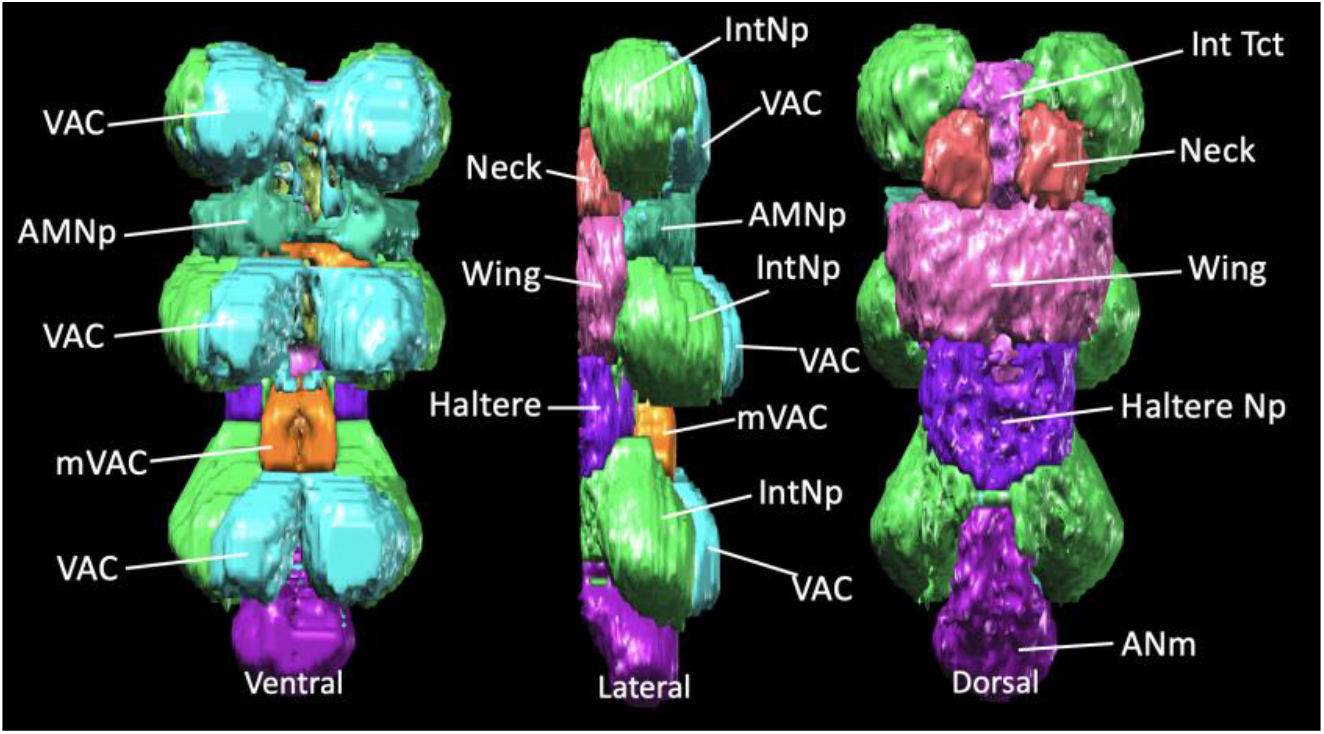
The major neuropils of the VNS. surface rendering of the major neuropils of the VNS showing structures from ventral, lateral and dorsal views (left to right).

#### Dorsal Medial Tract (DMT)

The **Dorsal Medial Tract** was previously called the Median Tracts of the Dorsal Cervical Connective (MTD) by Power, (1948) and Merritt and Murphey (1992); we propose DMT as an alternative to be more consistent with the rest of the tract nomenclature. The DMT is derived from the dorsal bundles in the cervical connective and extends posteriorly and bows laterally slightly, in the mesothoracic region, bending again medially, towards each other, at the narrowing between the meso- and metathoracic neuromeres (Figure 3 and 5). The tract turns laterally and posteriorly at the level of the haltere commissure and enters the metathoracic neuromere, where it forms collateral fibers that merge with the oblique tract of the metathoracic leg nerve (Power, 1948). The DMT contains the projections of a large number of descending neurons that innervate both the dorsal and ventral tectulum and the leg neuropils (Namiki et al., 2018).

**Figure 5.**
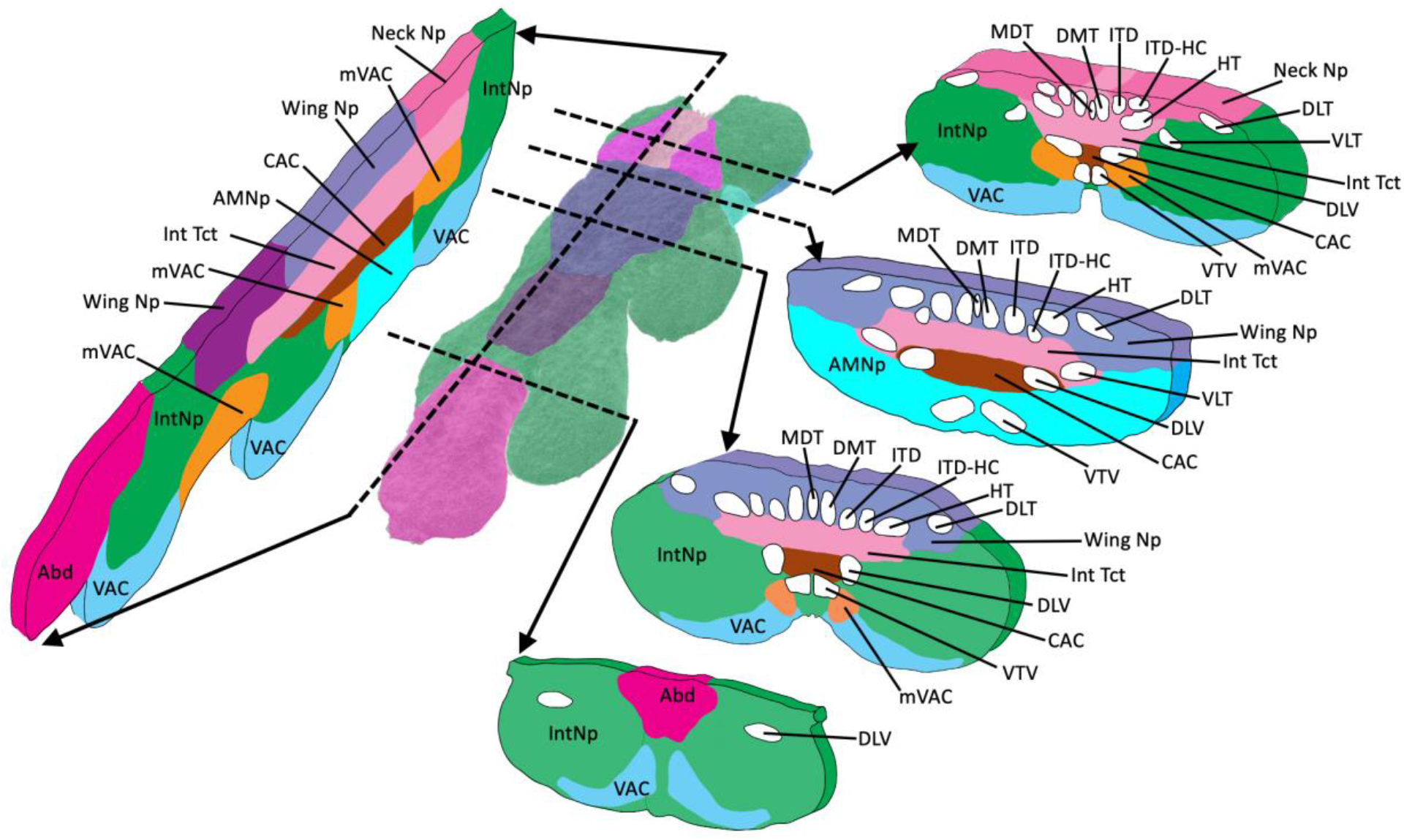
Schematic representation of the major neuropils and tracts of the VNS.

#### Dorsal Lateral Tract of Ventral Cervical Fasciculus (DLV)

The DLV derives from the ventral bundles of the cervical connective. It contains coarser fibers than the other tracts derived from the ventral bundles of the cervical connective (DLV, VLT and VTV). The tract broadens as it extends posteriorly, in a medial position and just ventral to the tectulum to terminate in the mesothoracic neuromere merging with the Oblique Tract (Figure 3 and 5).

#### Ventral Median Tract of Ventral Cervical Fasciculus (VTV)

VTV is the ventral-most tract in the VNS. It runs adjacent to the midline and derives from the ventral bundles of the cervical connective. The tract extends posteriorly either side of the midline just under the ventral most part of the lower tectulum (Figure 3 and 5) until they bend dorsally and terminates in the ventral-anterior region of the abdominal ganglion. A few fibers extend laterally in into the leg neuropil of each neuromere before the tract terminates in the abdominal ganglion (Power, 1948).

#### Median Dorsal Abdominal Tract (MDT)

**MDT** is the dorsal-most tract in the VNS, sits close to midline (Merritt and Murphey, 1992) and runs dorsally along the length of the tectulum (Boerner and Duch, 2010) past the haltere chiasma to terminate into the abdominal neuromeres (Power, 1948). It is the medial-most of the three small dorsal tracts which connects the thoracic and abdominal neuromeres (Figure 5).

## Discussion

With this nomenclature we address two primary issues required to create a clearer understanding of the VNS structure and to promote dialogue and data exchange amongst neuroscience researchers. The first was to establish a common anatomical framework to precisely define and describe, textually and spatially, the anatomical organization of the VNS. The second was to create a clear and consistent naming scheme for each anatomical entity. The detailed VNS map we provide is essential for integrating past and future work into a common space, thereby contributing to new lines of investigation. In addition, our effort will also inform researchers working with other insects, providing them with a template that can be adapted to their own model organism. Although the nomenclature developed in this project will serve as an initial standard, we acknowledge that to remain useful it must be maintained as a ‘living’ process and evolve as our understanding of the VNS structure and function grows. Future revisions and additions will be required, and this will be handled via the existing online system for posting anatomy ontology suggestions located at github.com/FlyBase/drosophila-anatomy-developmental-ontology/issues and maintained by VirtualFlyBrain.org.

Unlike the brain, the VNS in insects demonstrates significant diversity in its gross organization and structure (Niven et al., 2008). However, there is, a large anatomical literature for several insect groups that exhibit markedly different VNS structures (e.g. grasshoppers, crickets and moths) that often use the same terms as used for Drosophila. The differences amongst the VNSs of different insects are likely to be largely superficial and simply reflect the pattern of ganglionic fusion. Whilst this fusion does create some anatomical confusion, the basic pattern of tracts and commissures is preserved throughout the insects. Considering the conservation of lineages, tracts and commissures, insects do exhibit remarkably similar CNS structures despite the distortions imposed by ganglionic fusion. Consequently, it is important not only to have a consistent nomenclature to help work with Drosophila we also need to develop a nomenclature that can be used as broadly as possible across the insects to create a consistent cross species terminology. Whilst this would require some work to confirm homology rather than rely on inference from similar structure, extension of a consistent nomenclature to other insects would provide a framework to explore cross species homologies in the VNS and the deep evolutionary conservation of the nervous system.

## Acknowledgments

We thank Gerald M. Rubin and the staff of the Howard Hughes Medical Institute’s Janelia Research Campus for hosting our workshop. Special thanks to Wyatt Korff and the other members of the organising committee‡ for such a successful workshop. This work was initially supported in part by grants EP/F500385/1 and BB/F529254/1 for the University of Edinburgh School of Informatics Doctoral Training Centre in Neuroinformatics and Computational Neuroscience (http://www.anc.ed.ac.uk/dtc) from the UK Engineering and Physical Sciences Research Council (EPSRC), UK Biotechnology and Biological Sciences Research Council (BBSRC), and the UK Medical Research Council (MRC). Finally, by the Welcome Trust as part of the ‘Virtual Fly Brain: a global informatics hub for Drosophila neurobiology’ WT105023MA.

## Competing Interests

**Robert Court**

Informatics Forum, Edinburgh, EH8 9AB, Scotland

Contribution: Attended the workshop, contributed data, segmented data, managed curation of the data and wrote first draft of the manuscript

For Correspondence

Competing Interests: None

**Shigehiro Namiki**

Research Center for Advanced Science and Technology, The University of Tokyo, 4-6-1 Komaba, Meguro, Tokyo 153-8904, Japan

Howard Hughes Medical Institute, Janelia Research Campus, 19700 Helix Drive, Ashburn, VA 20147

Contribution: Attended the workshop, contributed data, segmented data and reviewed the manuscript

For Correspondence

Competing Interests: None

**J Douglas Armstrong**

School of Informatics, University of Edinburgh. 10 Crichton Street, Edinburgh, EH8 9AB

Contribution: Attended the workshop, contributed data and reviewed the manuscript

For Correspondence

Competing Interests: None

**Jana Börner**

Biological Sciences, College of Science John D. McArthur, Jupiter Campus, Jupiter, FL 33458

Contribution: Attended and participated in the workshop, contributed data and reviewed the manuscript

For Correspondence

Competing Interests: None

**Gwyneth Card**

Howard Hughes Medical Institute, Janelia Research Campus, 19700 Helix Drive, Ashburn, VA 20147

Contribution: Attended and participated in the workshop, contributed data and reviewed the manuscript

For Correspondence

Competing Interests:N

**Marta Costa**

Department of Zoology, University of Cambridge, Downing Street, Cambridge, CB2 3EJ UK

Contribution: Attended and participated in the workshop, curates the data and reviewed the manuscript

For Correspondence

Competing Interests: None

**Michael Dickinson**

Division of Biology and Bioengineering, California Institute of Technology, 1200 E. California Blvd., Pasadena CA 91125, USA

Contribution: Attended the workshop and reviewed the manuscript

For Correspondence

Competing Interests: None

**Carsten Duch**

Institute of Zoology, Johannes Gutenberg-University Mainz, 55128 Mainz

Contribution: Attended the workshop, contributed data and reviewed the manuscript

For Correspondence

Competing Interests: None

**Wyatt Korff**

Howard Hughes Medical Institute, Janelia Research Campus, 19700 Helix Drive, Ashburn, VA 20147

Contribution: Local Organizer, attended the workshop, provided data and reviewed the manuscript.

For Correspondence

Competing Interests: None

**Richard Mann**

Mortimer B. Zuckerman Mind Brain Behavior Institute, Columbia University,

Contribution: Attended and participated in the workshop and reviewed the manuscript

For Correspondence

Competing Interests: None

**David J Merritt**

School of Biological Sciences, The University of Queensland

Contribution: Contributed data and reviewed the manuscript

For Correspondence

Competing Interests: None

**Rod K Murphey**

Biological Sciences, College of Science John D. McArthur, Jupiter Campus, Jupiter, FL 33458

Contribution: Attended the workshop, contributed data and reviewed the manuscript

For Correspondence

Competing Interests: None

**Andrew Seeds**

University of Puerto Rico, Institute of Neurobiology, 201 Boulevard del Valle, San Juan, Puerto Rico 00901

Contribution: Attended and participated in the workshop and reviewed the manuscript

For Correspondence

Competing Interests: None

**Troy Shirangi**

Villanova University, 800 Lancaster Avenue, Villanova, PA 19085

Contribution: Attended the workshop and reviewed the manuscript

For Correspondence

Competing Interests: None

**Julie H. Simpson**

Department of Molecular, Cellular and Developmental Biology and Neuroscience Research Institute, University of California Santa Barbara, Santa Barbara CA 93106 USA

Contribution: Attended the workshop and reviewed the manuscript

For Correspondence

Competing Interests: None

**James W Truman**

Friday Harbor Laboratories, University of Washington, Friday Harbor, WA 98250, USA

Contribution: Attended the workshop, contributed data and reviewed the manuscript

For Correspondence

Competing Interests: None

**John Tuthill**

University of Washington,1705 N.E. Pacific Street, Seattle, WA 9819

Contribution: Attended the workshop and reviewed the manuscript

For Correspondence

Competing Interests: None

**Darren Williams**

Department of Developmental Neurobiology, King’s College London, New Hunt’s House, 4th Floor, Guy’s Hospital Campus, London SE1 1UL

Contribution: Attended the workshop, contributed data and reviewed the manuscript

For Correspondence

Competing Interests: None

**David Shepherd**

Address: School of Natural Sciences, Bangor University, Bangor, Gwynedd, UK LL57 2UW

Contribution: Convened and attended the workshop, contributed data and drafted final version of the manuscript the manuscript. Corresponding author.

For Correspondence

Competing Interests: None

